# Genomic determinants of pathogenicity in SARS-CoV-2 and other human coronaviruses

**DOI:** 10.1101/2020.04.05.026450

**Authors:** Ayal B. Gussow, Noam Auslander, Guilhem Faure, Yuri I. Wolf, Feng Zhang, Eugene V. Koonin

## Abstract

SARS-CoV-2 poses an immediate, major threat to public health across the globe. Here we report an in-depth molecular analysis to reconstruct the evolutionary origins of the enhanced pathogenicity of SARS-CoV-2 and other coronaviruses that are severe human pathogens. Using integrated comparative genomics and machine learning techniques, we identify key genomic features that differentiate SARS-CoV-2 and the viruses behind the two previous deadly coronavirus outbreaks, SARS-CoV and MERS-CoV, from less pathogenic coronaviruses. These features include enhancement of the nuclear localization signals in the nucleocapsid protein and distinct inserts in the spike glycoprotein that appear to be associated with high case fatality rate of these coronaviruses as well as the host switch from animals to humans. The identified features could be crucial elements of coronavirus pathogenicity and possible targets for diagnostics, prognostication and interventions.

---

The emergence of novel SARS-coronavirus-2 (SARS-CoV-2), which causes the respiratory disease COVID-19, triggered a global pandemic that has led to an unprecedented worldwide public health emergency^1^. Since it was first reported in December 2019 and as of April 7, 2020, SARS-CoV-2 has infected over a million individuals worldwide, and has led to an estimated 82,000 deaths, with its associated morbidity and mortality rates continuously rising^2^. SARS-CoV-2 is the seventh member of the *Coronaviridae* family known to infect humans^3^. SARS-CoV and MERS-CoV, two other members of this family, are the causative agents of recent outbreaks, accountable, respectively, for the severe acute respiratory syndrome (SARS, 2002-2003) and Middle East respiratory syndrome (MERS, began in 2012) outbreaks^3,4^, and are associated with high case fatality rates (CFR, 9% and 36%, respectively). The novel SARS-CoV-2 can also cause severe disease and is appreciably more infectious than SARS-CoV or MERS-CoV, but with a lower associated CFR^4^. By contrast, the other coronaviruses infecting humans, HCoV-HKU1, HCoV-NL63, HCoV-OC43, and HCoV-229E, are endemic and cause mild symptoms, accounting for 15-29% of common colds^3^. The three coronaviruses that can cause severe diseases (hereafter high-CFR CoV) originated in zoonotic transmissions from animal hosts to humans. SARS-CoV and MERS-CoV have bat reservoirs, and were transmitted to humans through intermediate hosts (likely civets and camels, respectively)^4^. Similarly, the closest known relative of SARS-CoV-2 is a bat coronavirus (Fig. 1a), but the specific route of transmission from bats to humans remains unclear. These repeated, independent zoonotic transmissions and the high associated pathogenicity call for an in-depth investigation of the genomic features that contribute to coronaviruses pathogenicity and transmission, to better understand the molecular mechanisms of the high-CFR CoV pathogenicity, and thus to be better prepared for any future coronavirus outbreaks, and potentially contribute to the development of interventions.

**Figure 1.**
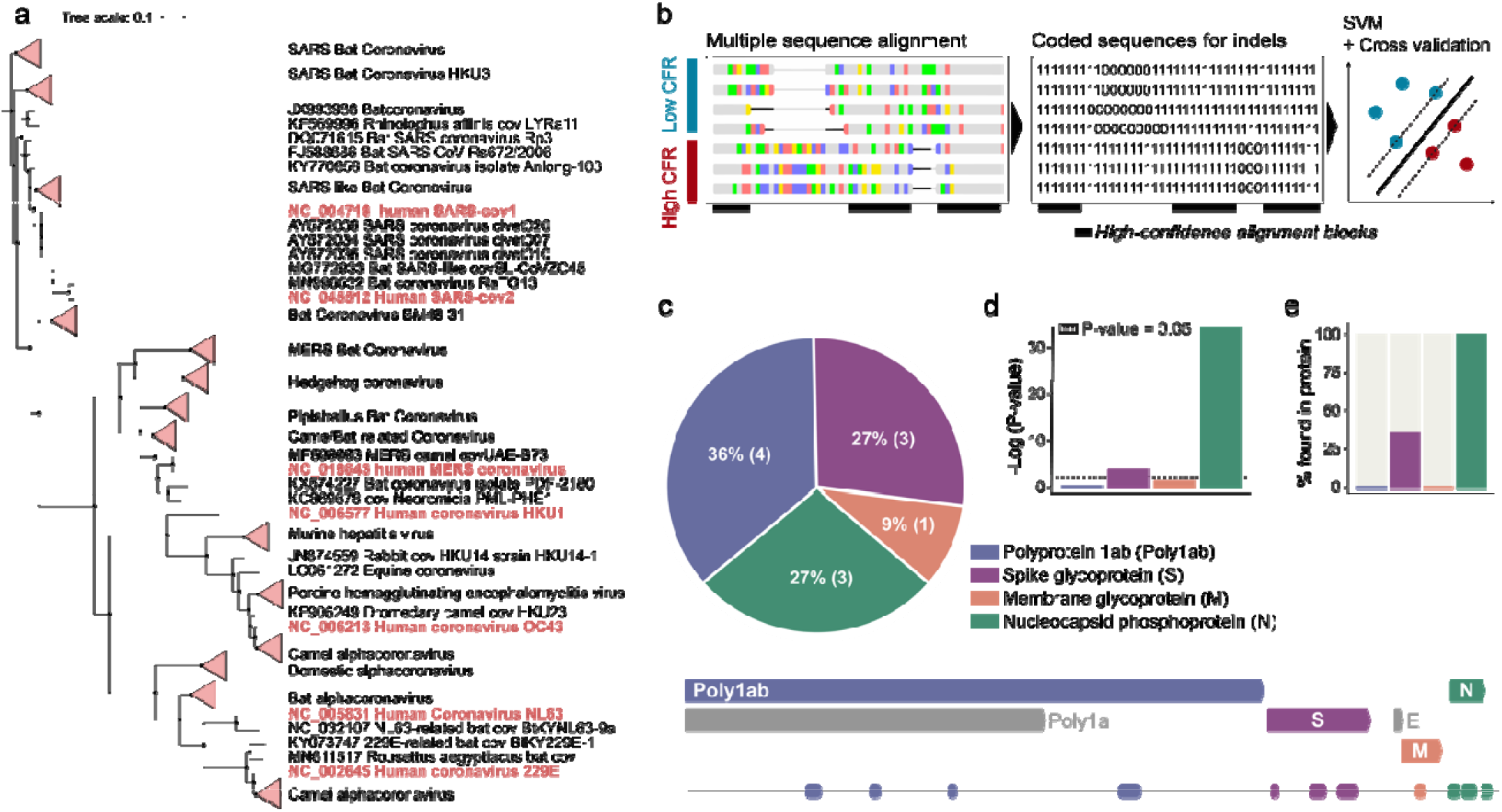
Searching coronavirus genomes for determinants of pathogenicity. **(a)** Phylogenetic tree of coronavirus species, based on the alignment of complete nucleotide sequences of virus genomes. **(b)** A schematic illustration of the pipeline applied for detection of genomic regions predictive of high-CFR strains. **(c)** Top panel: Pie chart showing the percentage of identified genomic determinants in each protein. Bottom panel: Map of SARS-CoV-2 genome with detected regions. **(d)** Bar plot showing the significance of the distribution of detected regions across each protein. **(e)** Percentage of detected predictive regions in each protein.

## Results

### Detection of diagnostic features of high-CFR coronaviruses

In this work, we developed an approach combining advanced machine learning methods with well-established genome comparison techniques, to identify the potential genomic determinants of pathogenicity of the high-CFR CoV strains (Fig. 1b). Coronaviruses have positive-sense RNA genomes consisting of 6 conserved proteins, along with additional strain-specific accessory proteins^5^. The conserved proteins are the polyproteins pp1a and pp1ab that encompass multiple protein domains involved in various aspects of coronavirus genome replication, spike glycoprotein (S), envelope (E), membrane glycoprotein (M) and nucleocapsid phosphoprotein (N, Fig. 1c). To detect potential genomic determinants of pathogenicity, we aligned the full genomes of all human coronaviruses (Supplementary File 1) and used support vector machines (SVM) to detect high-confidence genomic features that are predictive of the high CFR (see Methods for details). In total, our method identified 11 regions of nucleotide alignments that were reliably predictive of the high CFR of coronaviruses (Fig. 1c, Supplementary Table 1). Two proteins were significantly enriched with these predictive regions, the nucleocapsid protein and the spike glycoprotein (p-values: 4e-16 and 0.036, respectively, Fig. 1d). Only four of the diagnostic regions detected in the nucleotide alignment corresponded to observable differences in the protein alignments as well, with three located in the nucleocapsid protein and one in the spike protein (Fig. 1e).

### Enhancement of nuclear localization signals in the nucleocapsid protein

Exploring the regions identified within the nucleocapsid that predict the high CFR of CoV, we found that these deletions and insertions result in substantial enhancement of motifs that determine nuclear localization^6^, specifically, in high-CFR CoV (Supplementary Figure 1a). The deletions, insertions and substitutions in the N proteins of the high-CFR CoV map to two monopartite nuclear localization signals (NLS), one bipartite NLS and a nuclear export signal (NES, Fig. 2a,b). In the course of the evolution of coronaviruses, these nuclear localization and export signals grow markedly stronger in the clades that include the high-CFR viruses and their relatives from animals (primarily, bats), as demonstrated by the increasing positive charge of the amino acids comprising the NLS, a known marker of NLS strength^7^ (Fig. 2a). In the clade that includes SARS-CoV and SARS-CoV-2, the accumulation of positive charges was observed in the monopartite NLS, the bipartite NLS and the NES, whereas in the clade including MERS-CoV, positive charges accumulated primarily in the first of the two monopartite NLS (Fig. 2a, Supplementary Table 2). In all cases, the enhancement of these signals is a gradual, significant trend that accompanied coronavirus evolution concomitantly with the emergence of more pathogenic strains (empirical p-value < 0.001, Fig. 2a,c). The charge of the complete nucleocapsid protein gradually evolves towards greater positive values due, specifically, to the formation of the NLS, as demonstrated by sequence permutation analysis (Fig. 2a, see Methods for details), which implies a key role for these motifs in the function of the nucleocapsid including likely contribution to virus pathogenicity. The accumulation of positive charges resulting in strengthening of the NLS, which correlates with the growing CFR of coronaviruses (Fig. 2c), implies that the localization pattern of the nucleocapsid proteins of high-CFR strains differs from that of the low-CFR strains and might contribute the increased pathogenicity of the high-CFR strains. Localization of the nucleocapsid protein to the nuclei, and specifically, to the nucleoli, has been previously reported in coronaviruses^8^ and has been associated with increased pathogenicity in a porcine coronavirus model^9–11^. The presence of both NLS and NES raises an uncertainty as to the precise effect of these motifs on the nucleocapsid protein localization, and the reports are indeed somewhat contradictory^6,9,12^. Nevertheless, the striking extent of the changes in the NLS of the high-CFR strains (Fig. 2c) suggests that localization of the nucleocapsid protein is an important determinant of coronavirus pathogenicity.

**Figure 2.**
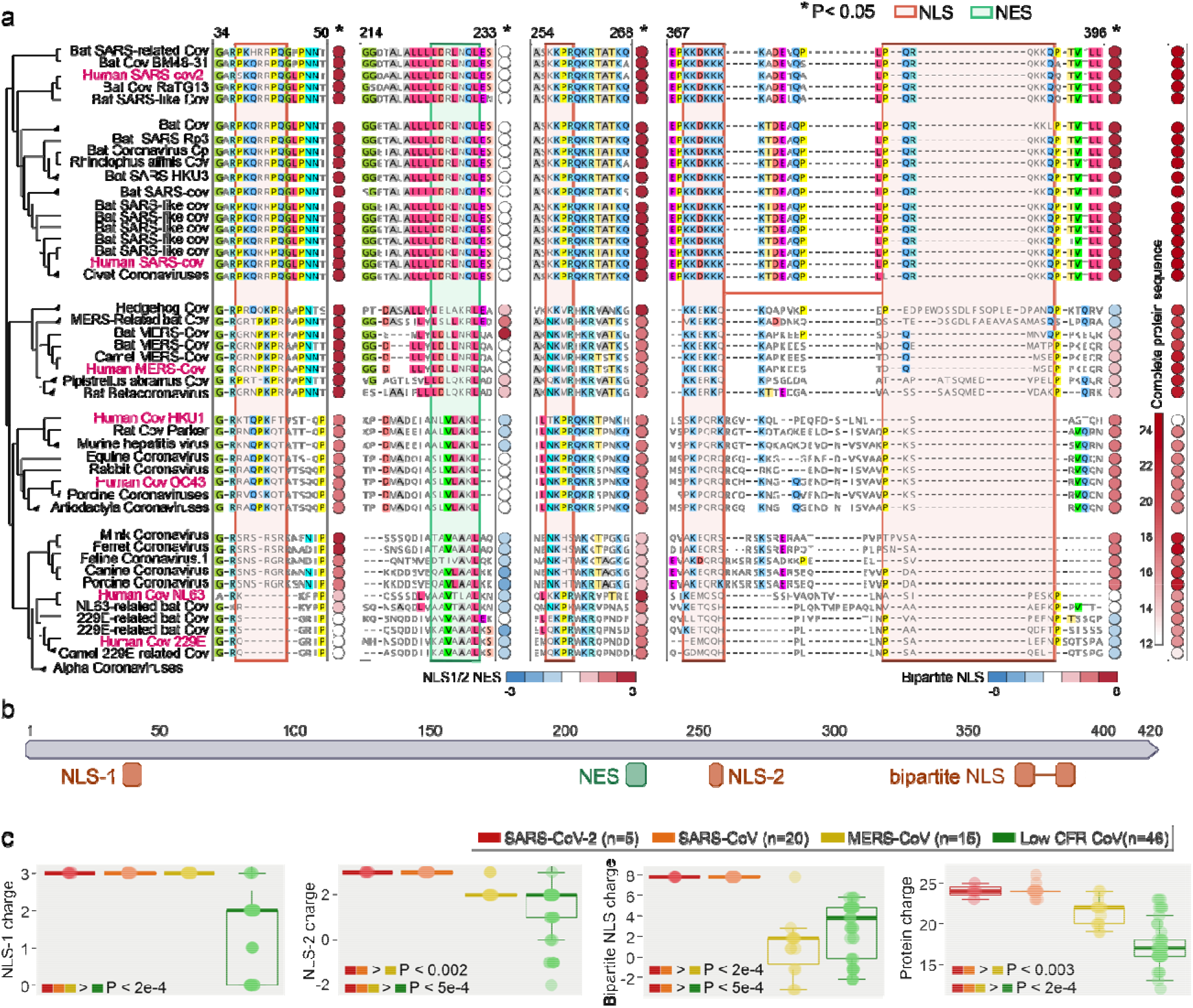
Putative determinants of coronavirus pathogenicity in the nucleoprotein and the spike protein. **(a)** Left panel: Phylogenetic tree and protein alignment of the nucleocapsid protein across coronavirus species. NLS and NES are outlined in red and green, respectively. The circle next to each signal sequence denotes peptide charge, with red denoting higher charge and blue denoting a lower charge. Right panel: The overall charge of each full protein sequence. **(b)** Map of SARS-CoV-2 nucleocapsid protein with relevant NLS (orange) and NES (green) motifs marked. **(c)** Boxplots displaying the charge of the three NLS motifs (left three panels) and that of the complete nucleocapsid protein (right panel) for SARS-CoV-2, SARS-CoV, MERS-CoV and low-CFR strains. The one-sided rank-sum P-values are shown when significant between any two groups, supporting a gradual increase of the charge.

### A unique insertion upstream of the heptad repeat region of the spike glycoprotein in high-CFR CoV

We next investigated the diagnostic feature identified within the spike glycoprotein. The SARS-CoV-2 spike protein binds ACE2, the host cell receptor of SARS-CoV-2^13^, with a 10-20 fold greater affinity compared to SARS-CoV, and contains a polybasic furin cleavage site resulting from a unique insert to SARS-CoV-2 that could enhance infectivity^4^. The spike protein consists of multiple domains^13^ (Fig 3a), including two heptad repeat regions that are crucial to infection^14^. During membrane fusion, the heptad repeats fold into a 6-helical bundle that forms the stable fusion core which facilitates the insertion of the hydrophobic fusion peptide into the host membrane and brings the viral and host membranes into proximity as required for fusion^15–17^. The spike protein fusion peptide is located upstream of the first heptad repeat^18,19^ (HR1), with a long connecting region between the fusion peptide and HR1 that adopts an α-helical structure. Our analysis revealed a 4 amino acid insertion in the connecting region in all high-CFR viruses but not in any of the low-CFR ones, with the MERS and SARS clades apparently acquiring this insertion independently as supported by the unrelated insert sequences (Fig 3b, Supplementary Figure 1b). The insertion increases the length and flexibility of the connecting region as confirmed by the examination of the spike glycoprotein structure of SARS-CoV (Fig. 3d), and therefore, is likely to affect the fusion process although the specific contribution of this insert to pathogenicity remains to be studied experimentally.

**Figure 3.**
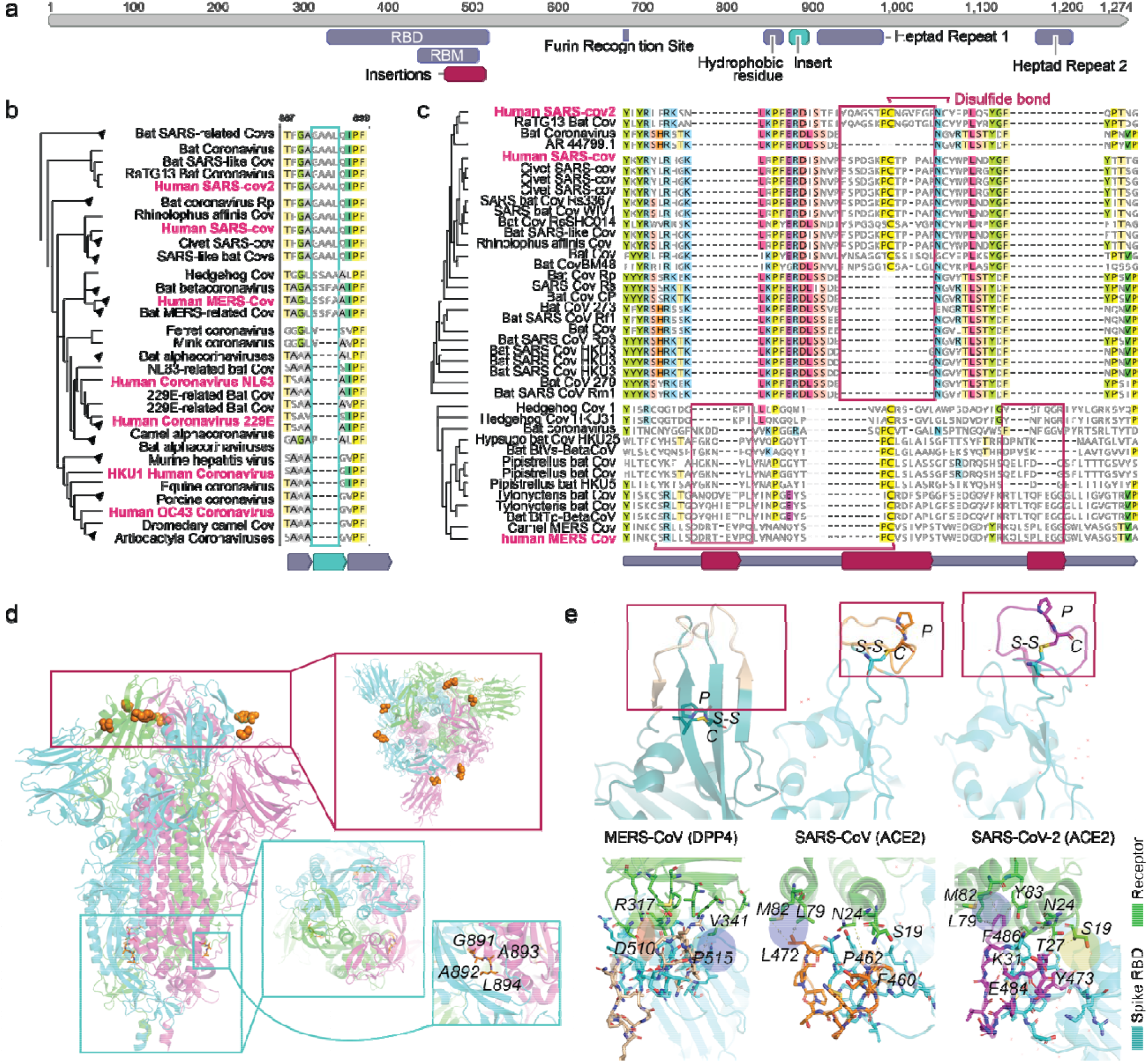
The signature inserts in the spike glycoproteins of the high-CFR CoV. **(a)** Map of SARS-CoV-2 spike protein with relevant protein regions and the features detected by the present analysis. The relevant regions include the receptor-binding domain (RBD), the receptor-binding motif (RBM), the furin recognition site, a hydrophobic residue preceding the first heptad repeat, and both heptad repeats. The two features detected by this analysis are the insertions in the RBM found in pathogenic strains before the zoonotic transmission to human, and the insertion in the high-CFR strains preceding the heptad repeat. **(b)** Phylogenetic tree and protein alignment of the spike protein insertion preceding the first heptad repeat. **(c)** Phylogenetic tree and protein alignment of the spike protein insertions in the RBM of high-CFR CoV (disulfide bonds are shown for human strains). **(d)** Structure of the SARS-CoV-2 spike glycoprotein trimer with the inserts mapped to the heptad repeat-containing domain and the receptor-binding domain. The top inset (red rectangle) shows the locations of the inserts in the RBM, designated as in (a), that are located within segments bordered by orange spheres (unresolved in the structure). The middle inset (blue rectangle) shows the GAAL insert upstream of the first heptad-repeat region. The bottom inset (green rectangle) shows a close view of the GAAL insert (in orange). **(e)** Top panel: Structures of the receptor-binding motifs of SARS-CoV, SARS-Cov-2 and MERS.

### Unique inserts associated with zoonotic jumps to humans in the spike glycoprotein

Finally, we sought to identify genomic features that might be associated with the repeated jumping of coronaviruses across the species barriers to humans, specifically, in the case of the high-CFR strains. To this end, we aligned the genomes of all coronaviruses from different hosts (Supplementary File 2) and selected, for each human-infecting strain of the high-CFR CoV, the closest non-human infecting relatives (see Methods for details). Within each such set of human high-CFR CoV and their animal ancestors, we searched for genomic insertions or deletions that occurred in the most proximal strains before the zoonotic jump to humans. This analysis identified independent insertions in each of the three groups of viruses all of which were located within the spike glycoprotein, specifically, in the receptor binding domain, within the subdomain that binds ACE2^13,20^ (the receptor binding motif, RBM) in the cases of SARS-CoV and SARS-CoV-2, and DPP4 in the case of MERS-CoV^21^ (Fig. 3c,d).

The insertions occur in slightly different locations within the RBM structure, with a single insert in the SARS clade and two distinct inserts in the MERS clade, and showed no sequence similarity between the three high-CFR groups, suggesting independent evolutionary events. The two insertions in MERS clade correspond to two loops connecting the distal β-strand of an extended β-sheet structure, whereas the insert in the SARS clade corresponds to a single long loop embedded within a short, unstable β-sheet. Despite the lack of an overall similarity, in each case, the inserted segments contained a proline-cysteine (PC) amino acid doublet (Fig. 3c,e and Supplementary Figure 2). In both high-CFR CoV clades, the cysteines in the insert form disulfide bonds with other cysteines in the RBM although the locations of the partner cysteines are different, i.e. proximal, within the loop containing the insert in SARS, and distal, within the RBM β-sheet in MERS (Fig. 3e and Supplementary Figure 2). These different locations of the disulfide bridges result in distinct RBM conformations (Fig. 3e) corresponding to the different receptor specificities in human cells. In both SARS and MERS, the insert directly interacts with the respective receptors but the specifics of the interactions differ, with a salt bridge (D510 with R317) and a hydrophobic interaction (P515 with V341) in the case of MERS, and a hydrophobic interaction patch in SARS-CoV and SARS-CoV-2 each, involving L472 and F486, respectively. The flexibility added to the RBM by these inserts could allow the spike to be more malleable in binding to a receptor, allowing for zoonotic transmission. Furthermore, the SARS inserts that are located within a short, unstable β-sheet provide more flexibility than the MERS inserts which are within longer, more rigid β-sheets, potentially contributing to MERS-CoV never fully adapting to human-to-human transmission^4^. The phenylalanine in SARS-CoV-2 provides a larger hydrophobic surface to interact with 3 residues of ACE2 (M82, L79 and Y83) whereas in SARS-CoV the leucine interacts with 2 residues (M82 and L79). The SARS-CoV-2 insert also engages in an extra charge interaction with ACE2 (E484 with K31) and multiple hydrogen bonds (Fig. 3d). The larger hydrophobic surface and the additional interactions could, in part, underlie the higher binding affinity of SARS-CoV-2 to ACE2 compared to SARS-CoV^13^. Thus, the independent insertions in the RBM of the spike protein are highly likely to contribute to or even enable the zoonotic transmission of the high-CFR CoV strains to humans and might also contribute to their high CFR.

The inserts are highlighted in wheat for MERS, orange for SARS-CoV, and purple for SARS-CoV-2, and the proline-cysteine doublets and disulfide bonds are shown. Bottom panel: Interactions between the inserts in the RBM of the spike glycoproteins of SARS-CoV, SARS-CoV-2 and MERS, and the corresponding human receptors. Residues shown with stick models are within a 5Å distance from the interacting residues in the inserts. The salt bridge is highlighted in red with a thick red border (in MERS), charge interaction is highlighted in red with a thin blue border (SARS2), and H-bond network is highlighted in yellow (Y473, T27, S19).

SARS-CoV-2 has led to the most devastating pandemic since the 1918 Spanish flu, prompting an urgent need to elucidate the evolutionary history and genomic features that led to the increased pathogenicity and rampant spread of this virus as well as those coronaviruses that caused previous deadly outbreaks. A better understanding of viral pathogenicity and zoonotic transmission is crucial for prediction and prevention of future outbreaks. Here, using an integrated approach that included machine-learning and comparative genomics, we identified three previously undetected likely determinants of pathogenicity and zoonotic transmission. The enhancement of the NLS in the high-CFR CoV nucleocapsids implies an important role of the subcellular localization of the nucleocapsid protein in the CoV pathogenicity. Strikingly, insertions in the spike protein appear to have been acquired independently by the SARS and MERS clades of the high-CFR CoV, in both the domain involved in virus-cell fusion and the domain mediating receptor recognition. These insertions, most likely, enhance the pathogenicity of high-CFR viruses and contribute to their ability to zoonotically transmit to humans. All these features are shared by the high-CFR CoV and their animal (in particular, bat) infecting relatives in the same clade, which is compatible with the possibility of future zoonotic transmission of additional highly pathogenic strains to humans. The predictions made through this analysis unveil critical features in the mechanism of SARS-CoV-2 virulence and evolutionary history, are amenable to straightforward experimental validation and could serve as predictors of strains pathogenic to humans.

## Methods

### Data

The complete nucleotide sequences of 3001 coronavirus genomes were obtained from NCBI (Supplementary File 2). Of these, 944 genomes belong to viruses that infect humans, including both viruses with low case fatality rates (CFR), NL63, 229E, OC43 and HKU1, and those with high CFR, namely, MERS, SARS-CoV-1 and SARS-CoV-2. The protein sequences that are encoded in the genomes of all human coronaviruses and closely related viruses from animals were obtained from NCBI, including the two polyproteins (1ab and 1a), spike glycoprotein, envelope, membrane glycoprotein, and nucleocapsid phosphoprotein.

### Identification of genomic determinants of high-CFR coronaviruses

To identify genomic determinants of coronaviruses associated with high CFR, comparative genome analysis was combined with machine learning techniques. First, the 944 human coronavirus genomes were aligned using Mafft^22^ v7.407. Then, we identified high confidence alignment blocks within the multiple sequence alignment (MSA), which were defined as regions longer than 15 bp, containing less than 10% of gaps in each position. We searched for regions containing deletions or insertions that separate high-CFR from low-CFR viruses and are surrounded by high confidence alignment blocks because these are most likely to contain relevant differences within conserved genomic regions. To this end, the aligned sequences were recoded such that each nucleotide was coded as ‘1’ and each gap as ‘0’. We then applied Support Vector Machines (using the Python library scikit-learn^23^ with a linear Kernel function) to a 5bp sliding window in the identified high-confidence alignment regions, with a leave-one-out cross validation (where all samples of one of the 7 coronaviruses were left out in each round of the cross validation, for a total of 7 rounds). Finally, we selected regions that predicted the high-CFR viruses with high confidence (greater than 80% accuracy) for further evaluation.

From the 11 regions identified, 4 were in the polyprotein 1AB, 3 in the spike glycoprotein, 1 in the membrane glycoprotein and 3 in the nucleocapsid phosphoprotein. To evaluate the significance of these finding, we computed a hyper-geometric enrichment P-value, using the sizes of the identified regions and the lengths of the coding regions of each protein within the MSA. We found that the nucleocapsid phosphoprotein was most enriched with genomic differences that predict CFR (P-value = 4e-16), followed by the Spike glycoprotein (P-value = 0.036) and that the polyprotein 1AB and membrane glycoprotein were not significantly enriched with such differences. We further examined the effects of this set of genomic differences on the resulting protein sequences, and found that only 4 of the 11 differences identified were reflected in the protein alignment, of which 3 occurred in the nucleocapsid phosphoprotein and one in the spike glycoprotein.

### Non-human proximal coronavirus strains

To compile a list of human and proximal non-human coronavirus strains, we first constructed a multiple sequence alignment of all 3001 collected strains using Mafft v7.407. From that alignment, we build a phylogenetic tree using FastTree^24^ 2.1.10 with the “-nt” parameter, and extracted the distances between leaves of each strain from each of the reference genomes of the 7 human coronaviruses (Supplementary File 3). We then extracted the proximal strains of each human coronavirus, which were within a distance less than 1.0 to one of the human coronaviruses. To obtain a unique set of strains, we removed highly similar strains by randomly sampling one strain from each group of strains with more than 98% pairwise sequence identity (the resulting strains are provided in Supplementary File 4).

### Amino acid charge calculations

To evaluate the strength of the identified NLS and NES motifs within the nucleocapsid phosphoprotein, we calculated the amino acid cumulative charge within the alignment region of each motif, and of the complete protein, for each of the selected human and proximal non-human coronavirus strains (Supplementary File 4). The charge of each region was evaluated by the number of positively charged amino acids (lysine and arginine) minus the number of negatively charged amino acids in that region (aspartic acid and glutamic acid). To evaluate the significance of the association between CFR and the charge of specific motifs within the nucleocapsid phosphoprotein, we first calculated the rank-sum P-value comparing the charges of regions in high-CFR versus low-CFR strains. Then, we applied a permutation test, by counting the fraction of similarly or more significant charge differentials values between high-CFR and low-CFR viruses within 1,000 randomly selected motifs with similar length from the alignment of the nucleocapsid phosphoprotein.

### Genomic determinants of the interspecies jump

To identify genomic determinants that discriminate high-CFR viruses that made the zoonotic transmission to humans, we used the nucleotide MSA of MERS, SARS-CoV-1 and SARS-CoV-2 and the selected proximal non-human coronaviruses of each of these.

We searched regions that maximize the following function:

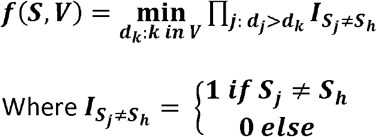

*S* is a position within the encoded MSA (‘1’ for a nucleotide and ‘0’ for a gap), and *V* is the set of strains selected for either MERS, SARS-CoV-1 or SARS-CoV-2. *d_k_* is the distance of non-human strain *k* from the human strain of group *V*, *S*_*j*_ is position *S* of non-human strain *j*, and *S*_*h*_ is position *S* of the human strain in group *V*.

Thus, this function aims to find, for each position, within each of the three groups of strains, the non-human strain *k* with the minimal distance from the human strain, such that all non-human strains that are more distant are more different than the human strain in that position (i.e. a genomic change that occurred as close as possible to the human strain). We searched for regions in which over 50% of the strains in the alignment differed from the human strain, and for which the differing strains were explicitly the most distant from human. We identified only one such location, across all three high-CFR virus groups.

### Structural analysis of the spike glycoproteins-receptor complexes

Crystal structures of the RBD spike glycoproteins of SARS-CoV (pdb: 2ajf^25^), SARS-CoV-2 (pdb: 6m0j^26^) and MERS (pdb: 4l72)^27^ complexed with their respective receptors, and the full cryoelectron microscopy structure of the SARS-CoV-2 (PDB: 6vxx)^28^ were downloaded from the Protein Data Bank^29^. Structural analyses including residues interactions and structural alignments were performed using the PyMOL computational framework^30^.

## Author contributions

EVK initiated the study; ABG, NA and YIW designed research; ABG, NA, and GF performed research; ABG, NA, GF, YIW, FZ and EVK analyzed the data; ABG, NA and EVK wrote the manuscript that was edited and approved by all authors.

## Acknowledgements

This research was supported by the Intramural Research Program of the National Library of Medicine at the NIH.

## Supplementary Information

Supplementary Table 1-2

Supplementary Figure 1-2

Supplementary datasets 1-4: ftp://ftp.ncbi.nih.gov/pub/wolf/_suppl/SARSpath20/

