## Supplementary tables and figures for "Genomic determinants of pathogenicity in SARS-CoV-2 and other human coronaviruses"

Supplementary Information

**Supplementary Tables**

| Alignment location | |  |  |  |
| --- | --- | --- | --- | --- |
| **start** | **end** | **gene** | Loc [SARS-Cov-2] | **type** |
| 7281 | 7291 | pp1ab | 3728 | indel |
| 10446 | 10451 | pp1ab | 6437 | indel |
| 14249 | 14249 | pp1ab | 9750 | indel |
| 25041 | 25108 | pp1ab | 19021 | indel |
| 29498 | 29498 | S (spike glycoprotein) | 22358 | indel |
| 32029 | 32040 | S (spike glycoprotein) | 24228 | indel |
| 32906 | 32927 | S (spike glycoprotein) | 25001 | indel |
| 36459 | 36462 | M (membrane glycoprotein) | 27001 | indel |
| 38798 | 38808 | N (nucleocapsid) | 29114 | indel |
| 38926 | 38941 | N (nucleocapsid) | 29235 | indel |
| 39356 | 39360 | N (nucleocapsid) | 29534 | indel |

**Supplementary Table 1.** The regions detected from the nucleotide genome alignment. The first two columns indicate the start and end positions of the detected region within the nucleotide genome alignment. Coordinates are inclusive. The third column indicates the gene that the region occurs in, the fourth column is the coordinate of the region in the SARS-CoV-2 reference, and the final column is the type of region detected.

|  | Motif1 (NLS1) | Motif2 (NES) | Motif3 (NLS2) | Motif4 (bipartite NLS) | Protein charge |
| --- | --- | --- | --- | --- | --- |
| YP_009724397 | 3 | 0 | 3 | 8 | 24 |
| QHR63308 | 3 | 0 | 3 | 8 | 24 |
| AVP78038 | 3 | 0 | 3 | 8 | 25 |
| ASO66816 | 3 | 0 | 3 | 8 | 24 |
| ABD75315 | 3 | 0 | 3 | 8 | 23 |
| ABG47067 | 3 | 0 | 3 | 8 | 24 |
| Q3I5I7 | 3 | 0 | 3 | 8 | 24 |
| AGC74175 | 3 | 0 | 3 | 8 | 24 |
| AHX37566 | 3 | 0 | 3 | 8 | 23 |
| AAZ41337 | 3 | 0 | 3 | 8 | 24 |
| Q3LZX4 | 3 | 0 | 3 | 8 | 24 |
| ADE34730 | 3 | 0 | 3 | 8 | 23 |
| AGC74169 | 3 | 0 | 3 | 8 | 24 |
| AAZ67039 | 3 | 0 | 3 | 8 | 24 |
| Q0Q468 | 3 | 0 | 3 | 8 | 26 |
| ACU31039 | 3 | 0 | 3 | 8 | 25 |
| ATO98190 | 3 | 0 | 3 | 8 | 24 |
| AGZ48815 | 3 | 0 | 3 | 8 | 24 |
| ARI44802 | 3 | 0 | 3 | 8 | 24 |
| AGZ48841 | 3 | 0 | 3 | 8 | 24 |
| NP_828858 | 3 | 0 | 3 | 8 | 24 |
| AAU04673 | 3 | 0 | 3 | 8 | 24 |
| AAU04658 | 3 | 0 | 3 | 8 | 24 |
| AAU04642 | 3 | 0 | 3 | 8 | 24 |
| APO40586 | 3 | 0 | 3 | 8 | 23 |
| YP_003858591 | 3 | 0 | 3 | 8 | 24 |
| QCC20721 | 3 | 1 | 3 | -3 | 21 |
| QGA70699 | 3 | 1 | 3 | -3 | 20 |
| AHY61344 | 3 | 2 | 2 | -1 | 20 |
| ASL68949 | 3 | 2 | 2 | -1 | 20 |
| YP_009361864 | 3 | 0 | 2 | 3 | 24 |
| AIG13103 | 3 | 0 | 2 | 2 | 23 |
| ASU90688 | 3 | 0 | 2 | 2 | 22 |
| YP_009047211 | 3 | 0 | 2 | 2 | 22 |
| YP_001039969 | 3 | 1 | 2 | 1 | 19 |
| QHA24694 | 3 | 1 | 2 | 1 | 19 |
| AWH65917 | 3 | 1 | 2 | 1 | 20 |
| ANA96046 | 3 | 1 | 2 | 2 | 22 |
| AIA62359 | 3 | 1 | 2 | 2 | 22 |
| AWH65884 | 3 | 1 | 2 | 2 | 22 |
| Q0Q4E6 | 3 | 1 | 2 | 2 | 22 |
| YP_001039960 | 3 | 1 | 2 | 2 | 22 |
| YP_173242 | 2 | -1 | 2 | 5 | 12 |
| YP_003029852 | 2 | -1 | 2 | 6 | 17 |
| YP_209238 | 2 | -1 | 2 | 6 | 18 |
| AAB86821 | 2 | -1 | 2 | 6 | 17 |
| NP_045302 | 2 | -1 | 2 | 6 | 17 |
| BAJ52884 | 2 | 0 | 2 | 5 | 17 |
| YP_005454249 | 2 | 0 | 2 | 5 | 16 |
| YP_009555245 | 2 | 0 | 2 | 5 | 16 |
| AAY68302 | 2 | 0 | 2 | 5 | 16 |
| ARC95207 | 2 | 0 | 2 | 5 | 16 |
| BAF75637 | 2 | 0 | 2 | 5 | 16 |
| ACX46856 | 2 | 0 | 2 | 5 | 16 |
| P10527 | 2 | 0 | 2 | 5 | 16 |
| ACJ66950 | 2 | 0 | 2 | 5 | 16 |
| ABG89284 | 2 | 0 | 2 | 5 | 16 |
| AHN64778 | 2 | 0 | 2 | 5 | 17 |
| ACB30189 | 2 | 0 | 2 | 5 | 16 |
| ABI94003 | 2 | 0 | 2 | 5 | 16 |
| AZU96330 | 2 | 0 | 2 | 5 | 16 |
| YP_009019186 | 3 | -1 | 1 | 3 | 22 |
| YP_009256201 | 3 | -1 | 1 | 3 | 23 |
| AFH58015 | 2 | -2 | 1 | 5 | 18 |
| AFG19745 | 2 | -2 | 1 | 4 | 21 |
| AMB66493 | 2 | -2 | 1 | 5 | 22 |
| YP_003771 | 1 | -1 | 3 | 0 | 16 |
| YP_009328939 | 1 | -1 | 2 | 0 | 18 |
| QHA24669 | 0 | -2 | 1 | -1 | 17 |
| APD51503 | 0 | -1 | 2 | -1 | 16 |
| NP_073556 | 0 | -1 | 2 | -2 | 16 |
| YP_009194643 | 0 | -1 | 2 | -2 | 13 |
| AOI28262 | 0 | -1 | 2 | -2 | 14 |
| ATI09441 | 0 | -1 | 2 | -2 | 14 |
| ASL24656 | 1 | 0 | -1 | 4 | 23 |
| AFE85966 | 1 | -2 | -2 | 2 | 20 |
| YP_001351688 | 1 | 0 | -1 | 4 | 20 |
| QCX35174 | 0 | 0 | -1 | 0 | 18 |
| YP_009201734 | 0 | -1 | 0 | 3 | 14 |
| QHA24675 | 0 | -1 | -1 | 3 | 23 |
| YP_001718609 | 0 | -2 | -1 | 3 | 17 |
| QHA24714 | 0 | 0 | -1 | -1 | 19 |
| YP_006908646 | 0 | 0 | -1 | -1 | 18 |
| QCX35164 | 0 | -1 | 1 | 0 | 16 |
| AIA62275 | 0 | 0 | 0 | -1 | 17 |
| YP_009199794 | 0 | 0 | 0 | -1 | 16 |

**Supplementary Table 2.** NLS and NES motifs charges and nucleocapsid protein charge of coronavirus strains considered.

**Supplementary Figures**

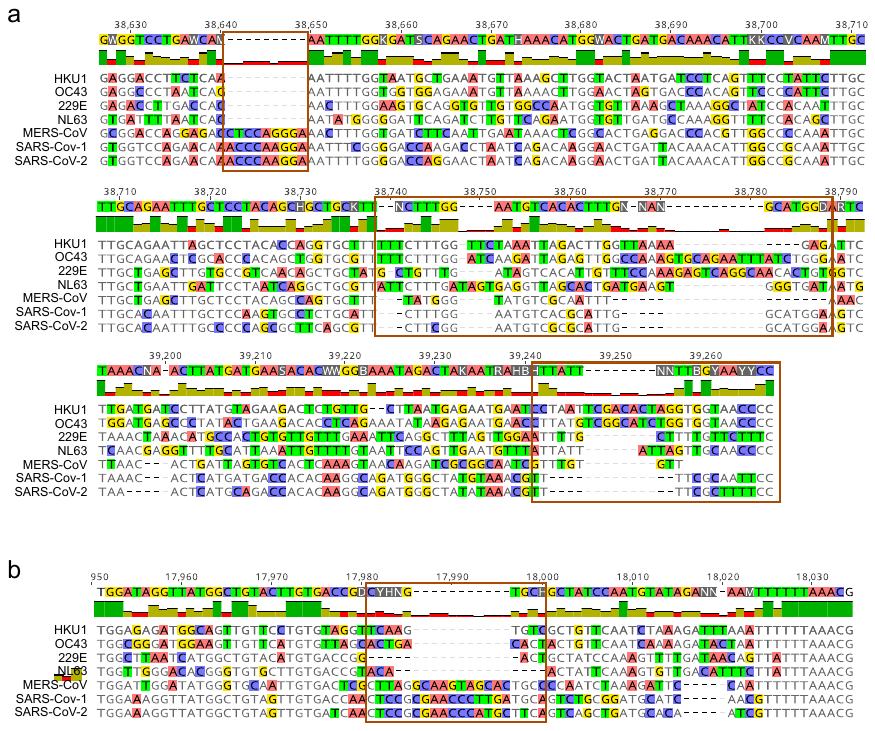

**Supplementary Figure 1. (a)** The locations detected within the nucleocapsid protein from the nucleotide genome alignment. **(b)** The location detected within the Spike protein from the nucleotide genome alignment.

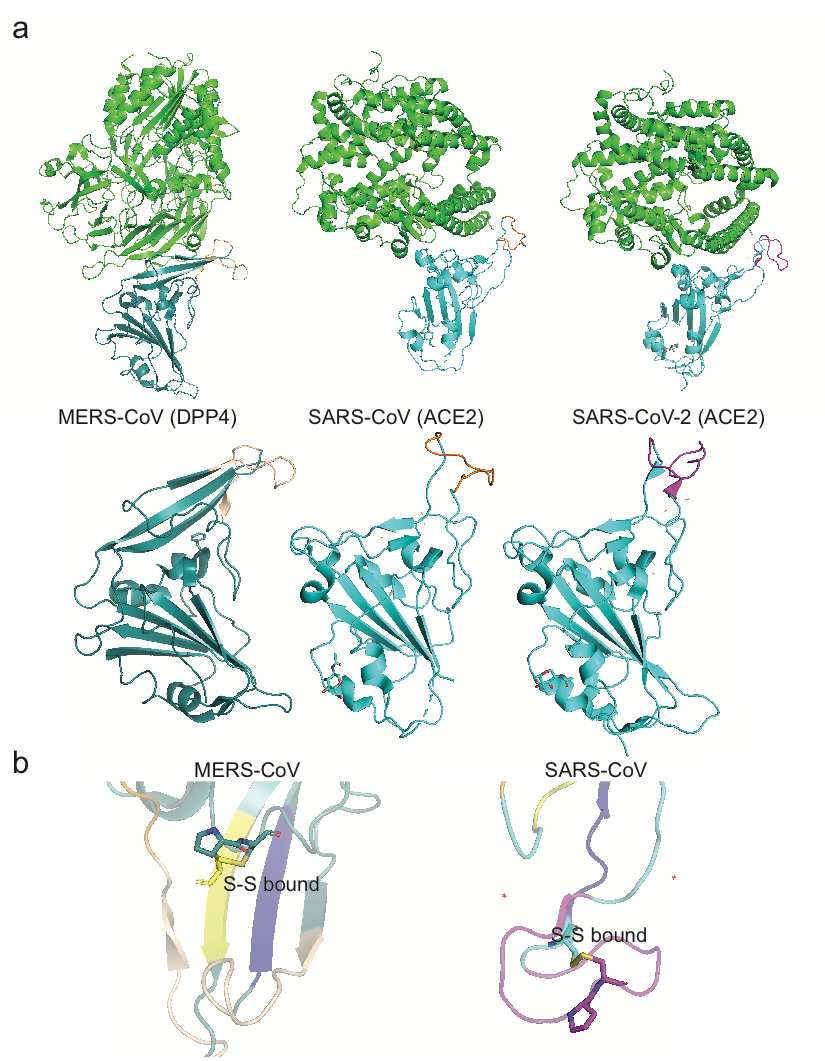

**Supplementary Figure 2. (a)** Complete structures of the receptor-binding motifs of SARS-CoV, SARS-Cov-2 and MERS (blue) and the human receptor that they bind to (green) **(b)** Close-up of the Structures of the receptor-binding motifs of SARS-CoVs and MERS-Cov. The inserts are highlighted, and the disulfide bonds are shown.
